# Quantifying brain state transition cost via Schrödinger bridge

**DOI:** 10.1101/2021.05.24.445394

**Authors:** Genji Kawakita, Shunsuke Kamiya, Shuntaro Sasai, Jun Kitazono, Masafumi Oizumi

## Abstract

Quantifying brain state transition cost is a fundamental problem in systems neuroscience. Previous studies utilized network control theory to measure the cost by considering a neural system as a deterministic dynamical system. However, this approach does not capture the stochasticity of neural systems, which is important for accurately quantifying brain state transition cost. Here, we propose a novel framework based on optimal control in stochastic systems. In our framework, we quantify the transition cost as the Kullback-Leibler divergence from an uncontrolled transition path to the optimally controlled path, which is known as Schrödinger bridge. To test its utility, we applied this framework to functional magnetic resonance imaging data from the Human Connectome Project and computed the brain state transition cost in cognitive tasks. We demonstrate correspondence between brain state transition cost and the difficulty of tasks. The results suggest that our framework provides a general theoretical tool for investigating cognitive functions from the viewpoint of transition cost.

**Author Summary:** In our daily lives, we perform numerous tasks with different kinds and levels of cognitive demand. To successfully perform these tasks, the brain needs to modulate its spontaneous activity to reach an appropriate state for each task. Previous studies utilized optimal control in deterministic systems to measure cost for brain state transition. However, there has not been a unified framework for quantifying brain state transition cost that takes account of stochasticity of neural activities. Here, we propose a novel framework for measuring brain state transition cost, utilizing the idea of optimal control in stochastic systems. We assessed the utility of our framework for quantifying the cost of transitioning between various cognitive tasks. Our framework can be applied to very diverse settings due to its generality.

## Introduction

The brain is considered a dynamical system that flexibly transitions through various states^1–3^. Depending on the properties of a dynamical system (e.g., the biophysical properties of neurons and the connectivity between neurons), some transitions are difficult to realize. Thus, characterizing the dynamical properties of brain state transition would be important for understanding various brain functions^4^, including decision-making^5^, motor control^6^, and working memory^7^, with potential applications in the diagnosis and clinical treatment of disease^8–10^. To date, however, no unified framework for quantifying brain state transition cost from brain activity data has been available.

One promising framework for quantifying brain state transition cost is the network control-theoretic framework^11^;see also^12,13^ for some limitations. Control theory provides useful perspectives for measuring the cost required for controlling a dynamical system to reach a desirable state. Considering the brain as a dynamical system, control-theoretic approaches enable us to quantify the cost of transitioning to a brain state that produces desirable behavior. Recently, the network control theoretic framework was proposed for study of the control property of the brain by viewing the brain as a networked dynamical system^14–17^. Although the framework provides an important new perspective for fundamentally understanding brain state transition, it has two major limitations. First, it does not capture the stochasticity of brain activity, which is ubiquitous in brain activity and is essential for accurately describing brain dynamics^18–20^. Disregarding stochasticity may result in an inaccurate estimation of transition cost. Second, the model obtained from structural connectivity, which is static over time, may not be able to capture change in the functional dynamics of the brain^4^, such as while performing tasks, for instance. Moreover, it is difficult to model even the resting state dynamics from structural connectivity^21^. Recently, alternative models using functional and effective connectivity has been proposed^22,23^, but these models still do not capture stochasticity. Thus, no unified framework able to take account of the key properties of brain dynamics is available.

Here, by employing control-theoretic approaches, we propose a novel framework for measuring brain state transition cost that can account for stochasticity. In our framework, we consider transition from a probability distribution of brain states to another distribution, rather than a transition from one brain state to another brain state (i.e. a point-to-point transition in a state-space) in contrast to a previous work^17^ utilizing network control theory. To transition from an initial distribution to a target distribution, the brain needs to modulate (control) its baseline transition probability. Although there are many possible ways to reach the target distribution, in this study we consider the optimally controlled path only and estimate the lower bound of brain state transition cost. We propose defining the minimum brain state transition cost as the Kullback-Leibler (KL) divergence from the baseline uncontrolled path to the optimally controlled path, i.e., the closest path to the original path, with the fixed initial and target distributions. The problem of finding the closest path to the original path connecting the initial and target distribution is known as Schrödinger bridge problem^24^, which has been studied in the fields of stochastic process and optimal transport^25–29^.

Here, as proof of concept, we apply the proposed framework based on Schrödinger bridge problem to evaluate the cost of task switching^30^, an executive function for moving from one task to another. Specifically, we address two questions. First, is the cost of transitioning to a more difficult task larger? A previous study^31^ reported that performing effortful tasks drives larger reconfiguration of functional brain networks. We therefore hypothesized that transitioning to a more difficult task required a larger cost. Second, is the brain state transition cost asymmetric? Specifically, is the transition cost from an easier task to a more difficult task larger than the cost accompanying the reverse transition?

To address these questions, we apply our framework to functional magnetic resonance imaging (fMRI) data from the Human Connectome Project (HCP)^32^. We use fMRI data of *n* = 937 subjects in the resting state and in seven cognitive tasks. After preprocessing and parcellation, we computed the probability distributions of coarse-grain brain activity patterns for the rest and cognitive tasks^17,33^. We then calculated the brain state transition cost by finding the Schrödinger bridge, i.e., the optimally controlled path^34^ between the initial and target distributions of brain states. We found that the transition cost to a more difficult task carried a larger transition cost. We also observed that the transition cost between an easy and a difficult task is asymmetric.

Overall, our findings provide a new perspective on the investigation of brain state transition, which may facilitate our understanding of cognitive functions.

## Results

### Quantification of brain state transition cost from Schrödinger bridge problem

In this study, we propose a novel framework to quantify state transition cost in a stochastic neural system, building on the formulation of Schrödinger bridge problem^25^. We consider brain state transition to be the transition from an initial probability distribution of brain states to a target probability distribution. In order to reach the target probability distribution, the brain is assumed to follow some controlled paths. Although there are many possible paths that bridge the initial and target probability distributions, we look for the optimally controlled path which minimizes the Kullback-Leibler (KL) divergence from an uncontrolled to a controlled path. Here, we define brain state transition cost as the minimum KL divergence from an uncontrolled to a controlled path that bridges the initial and target probability distributions (Fig. 1).

**Figure 1:**
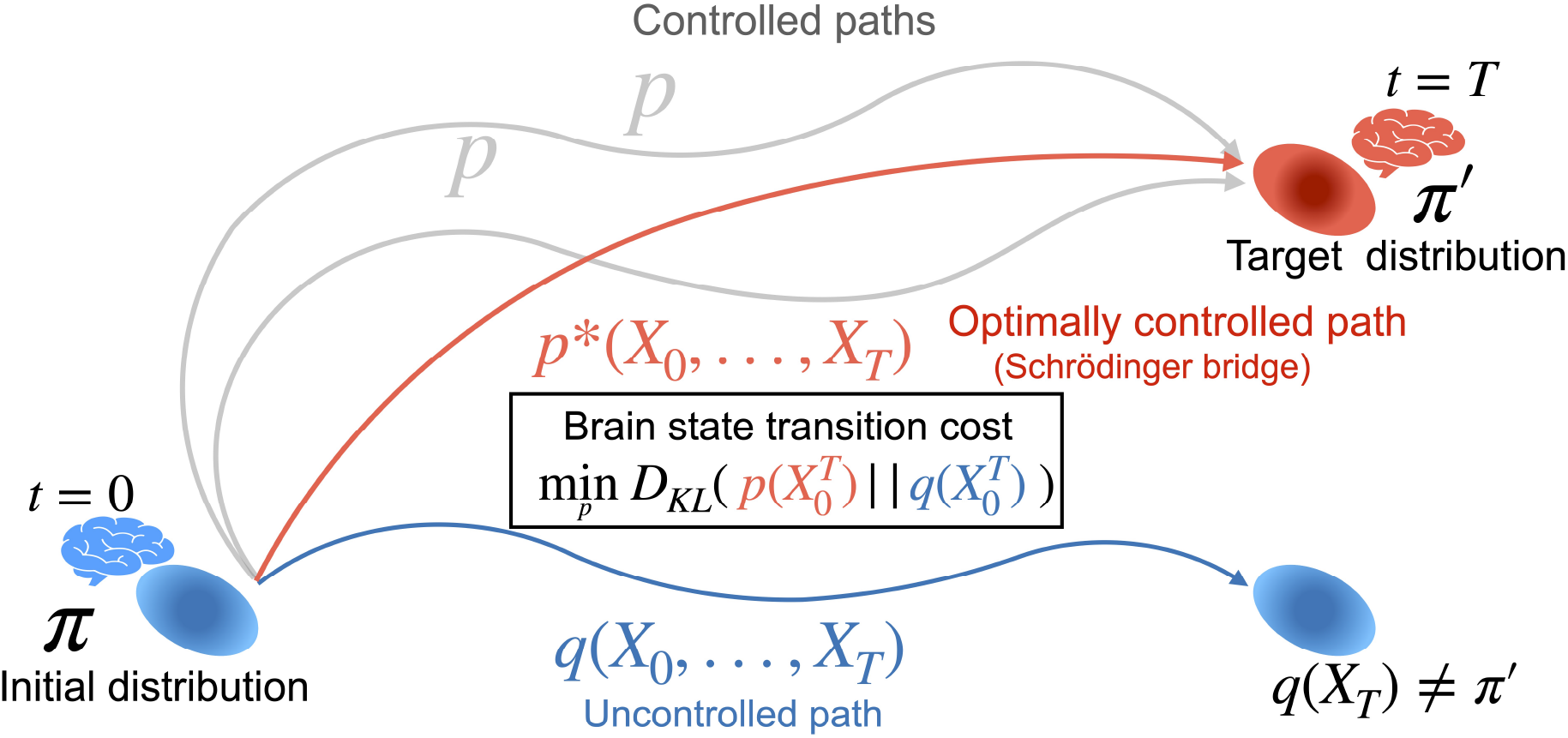
Schematic of brain state transition reframed as Schrödinger bridge problem. We consider brain state transition as transition from an initial probability distribution of brain states, *π*, to a target probability distribution, *π*′. The brain follows an uncontrolled baseline path, 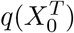, which does not lead to the target distribution but to *q*(*X_T_*) ≠ *π*′, where *q*(*X_T_*) represents the probability distribution at *t* =, following the uncontrolled path. In order to reach the target distribution, the brain needs to follow some controlled path, 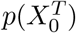. The brain state transition cost is defined as the minimum Kullback-Leibler divergence between the controlled and uncontrolled paths 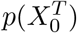 and 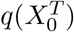, respectively. Optimally controlled path, 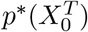 is equivalent to Schrodinger’s bridge.

We can mathematically formulate brain state transition as follows. Let *X_t_* be a random variable corresponding to a coarse-grained brain state at time *t*. We consider each brain state as a discrete number included in the finite set 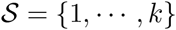, where *k* is the number of brain states. For instance, *X_t_* = *i* means that the brain is at the state *i* at time *t*. In this study, we used *k*-means clustering algorithm to obtain these coarse-grained brain states from high-dimensional brain activity data as described later. Then, let (*X*_0_,..., *X_T_*) be a time series of brain states that forms a first-order Markov chain. We introduce a simplified notation for expressing the time series as 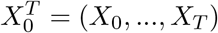 where subscript 0 represents the starting time point and superscript *T* represents the ending time point. We denote the joint probability distribution of the random variables by 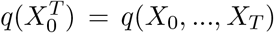, which can be expressed by using Markov property as follows:

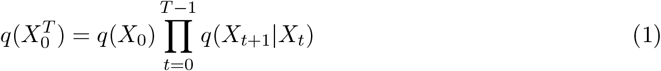

Here, we consider a problem of controlling the distribution of brain states to a target distribution, *π*′, at *t* = *T* starting from an initial distribution, *π*, at *t* = 0. The initial distribution, *π*, is the same as *q*(*X*_0_), but the target distribution *π*′ is different from *q*(*X_T_*), i.e., the target distribution, *π*′, cannot be reached by the original dynamics, 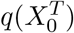. Thus, the transition probability of the brain needs to be modulated by some control input to the system. Although we do not explicitly model control input in this study because we do not model the dynamics of the system (see^29^ for the model of linear systems with control input), we assume that some control input is implicitly applied to modulate the original dynamics. In this context of controlling the dynamics, we call the original dynamics, 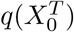, the “uncontrolled path”. In contrast with the uncontrolled path, 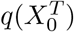, we call a controlled dynamics, 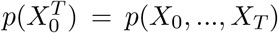, a “controlled path”, which satisfies the endpoint constraints, *p*(*X*_0_) = *π* and *p*(*X_T_*) = *π*′. While there are many possible controlled paths that satisfy the marginal conditions, we consider the problem of finding the optimally controlled path that minimizes some control cost. In this study, we define the control cost as the Kullback-Leibler (KL) divergence between the uncontrolled path and a controlled path^25,29^,

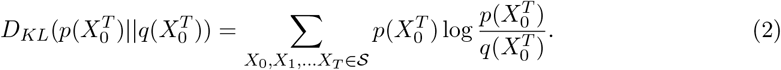

Intuitively, KL divergence measures the difference between two probability distributions. If KL divergence between 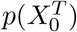 and 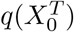 is zero, then we can tell that two paths are equivalent, i.e. the system does not change but stays the same. If the KL divergence takes nonzero values, it indicates that the system follows a different path from the uncontrolled path. Using the KL divergence as a transition cost is reasonable since the degree of the KL divergence should reflect how different a controlled path is from the uncontrolled path. Here, we define the optimally controlled path, 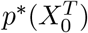, as the minimizer of the KL divergence as shown bellow

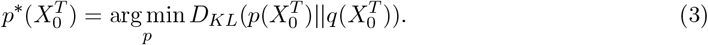

In words, the optimally controlled path, 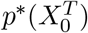, is the “closest” to the uncontrolled path, 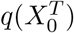, in terms of KL divergence. Then, by using the optimally controlled path, we define the minimum control cost 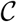 as

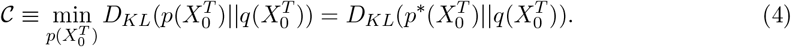

In this study, we propose to use the minimum control cost for quantifying the brain state transition cost. The problem of finding the optimally controlled path from an initial to a target distribution is known to be mathematically equivalent with Schrödinger bridge problem, the problem of finding the most likely path linking the initial and target distribution given the transition probability distribution of the system^25,29^.

To solve the minimization problem, we first decompose the KL divergence into two terms, both of which are also KL divergences.

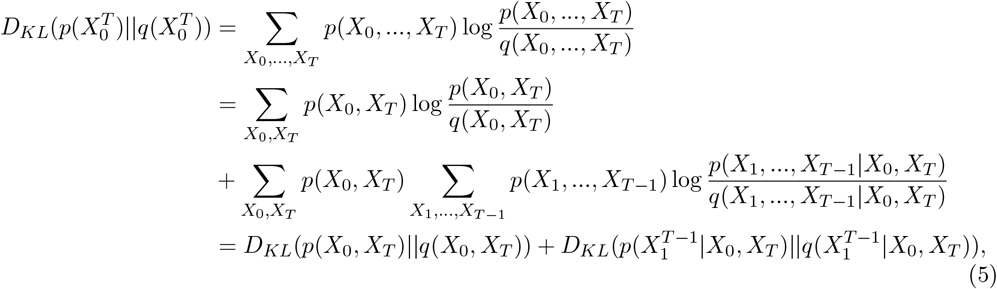

where 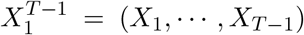. The two terms are both non-negative and we can separately minimize the two terms. The minimum of the second term is obviously 0 when 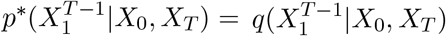. Then, the minimization problem of finding the whole controlled path is reduced to the problem of finding the optimally controlled joint distribution of the end points, *p*(*X*_0_,*X_T_*)^25^,

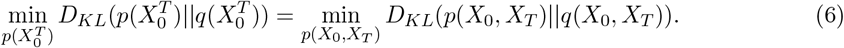

By introducing the useful notation *p*(*X*_0_, *X_T_*) = *P*, and *q*(*X*_0_, *X_T_*) = *Q*, where *P_ij_* = *p*(*X*_0_ = *i,X_T_* = *j*) and *Q_ij_* = *q*(*X*_0_ = *i, X_T_* = *j*) for the ease of computation, we can rewrite the KL divergence as

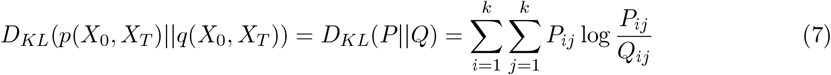

With these new notations, we can restate the original minimization problem (Eq. (3)) as

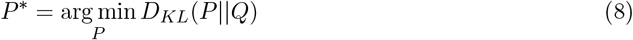

with the following constraints,

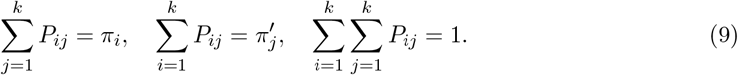

To further clarify the mathematical property of the optimization problem, we rewrite the control cost as follows.

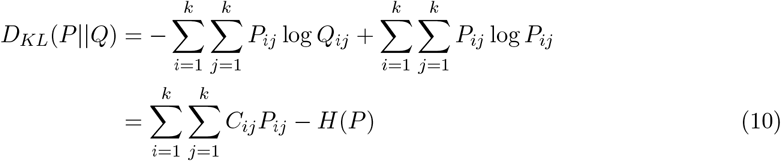

where *H*(*P*) is the entropy of the joint endpoint distribution, *P*. By rewriting the cost in this way, we can regard the problem of minimizing the KL divergence as the entropy regularized optimal transport problem with the transportation cost matrix, *C_ij_* = − log *Q_ij_*^35,36^; see e.g.^28,37,38^ for the connection between Schrödinger bridge problem and optimal transport problem. The existence and uniqueness of the optimal solution, *P*\*, is guaranteed because this is a strongly convex optimization _problem_^35,36^.

Here, we explicitly find the solution of the optimization problem by the method of Lagrange multipliers. Let 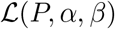 be the Lagrangian of Eq. (10) with Lagrange multipliers, *α_i_* and *β_j_*.

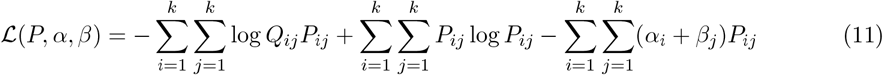

Differentiating the Eq. (11) with respect to *P_ij_* yields.

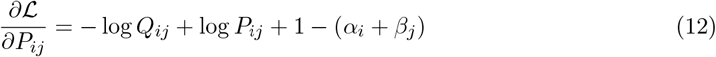

By setting the partial derivative to 0, we obtain the following optimal solution.

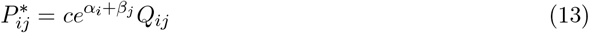

where *c* is the normalization constant,

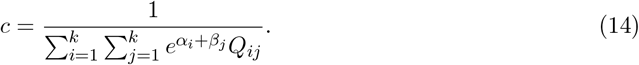

We determine the Lagrange multipliers, *α_i_* and *β_j_* by the constraints in Eq. (9),

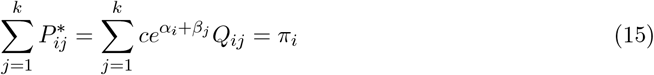

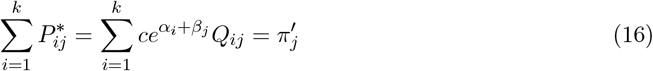

With some manipulation of the above equations, we obtain

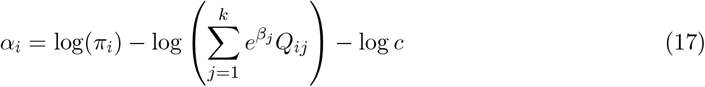

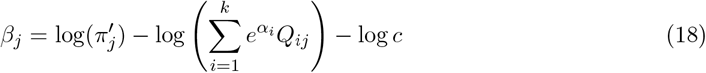

These Lagrange multipliers can be numerically determined by iteratively updating *α_i_* and *β_j_* according to the above equations starting from arbitrary initial values. This algorithm is known as Sinkhorn algorithm^35,39^. The implementation of the algorithm is available at https://github.com/oizumi-lab/SB_toolbox.

### Quantification of brain state transition cost in fMRI data

To test the utility of our proposed method, we applied the Schrödinger bridge-based framework to real fMRI data. We used resting-state fMRI and task-fMRI (emotion, gambling, language, motor, relational, social, and working memory) from the Human Connectome Project (HCP)^32^. We first performed preprocessing of the BOLD signals and parceled them into 100 regions^40^. As shown in Fig. 2, we concatenated the preprocessed data of all subjects for all the tasks to obtain *M* × *N* time series data, where *M* is the number of parcels (100), and *N* is the total time frames of the concatenated data. We consider a point in *M* = 100 dimensional space as the activity of the whole brain at a particular time frame. In total, there are *N* points in this high dimensional space. We applied the *k*-means clustering algorithm to classify *N* points into *k* coarse-grained states. In this section, we show only the results when we set the number of coarse-grained states to *k* = 8 (see Supporting information for the results with different numbers of coarse-grained states).

**Figure 2:**
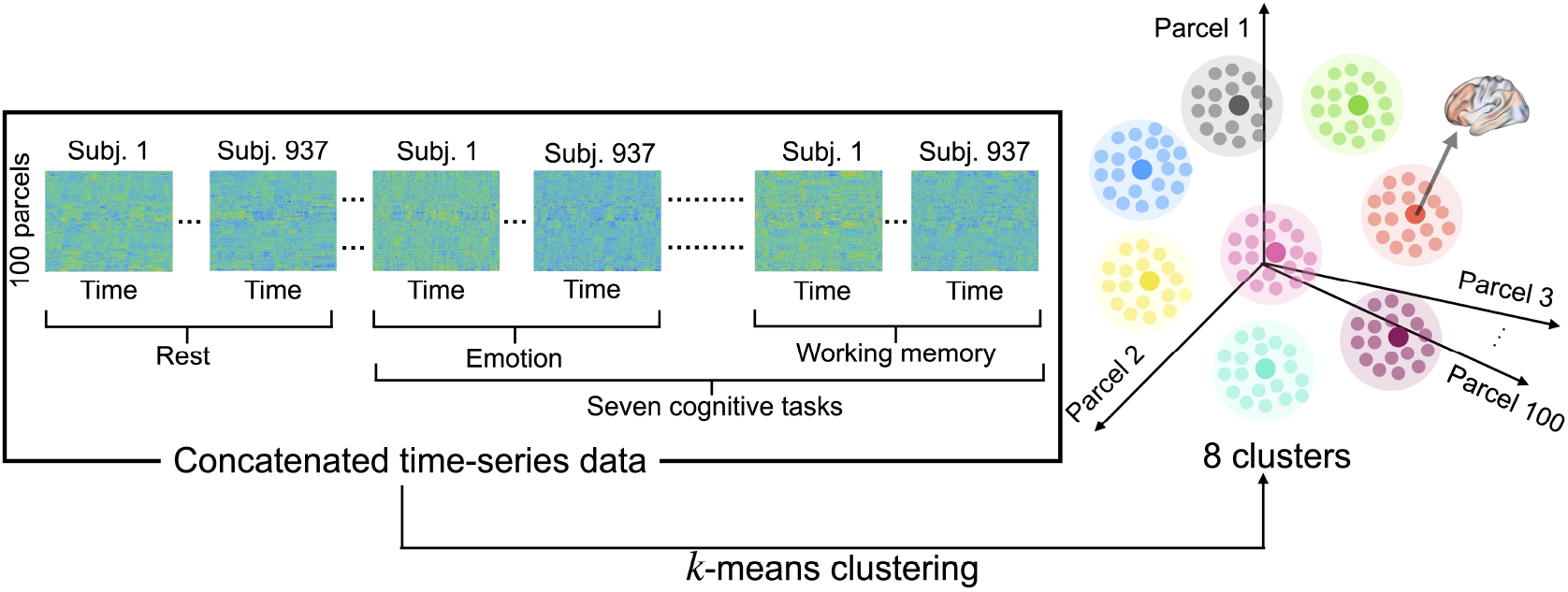
Clustering fMRI data. After preprocessing raw fMRI data, we concatenated the preprocessed data of all subjects for all the tasks. We then used *k*-means clustering to group similar brain activity patterns into eight coarse-grained brain states. Each point in the 100 dimensional state space corresponds to the activity of the whole brain at a particular time frame (see S5 in the Supporting information for brain maps of the centroids of the eight clusters).

To compute the brain state transition cost Eq. (4), we need to obtain initial and target distributions as well as the joint probability distribution for the uncontrolled path. Here, we assume that an uncontrolled path in the brain is the resting state transition probability. To determine the joint probability distribution for the uncontrolled path, we need to set the value of *T*, the time steps to reach the target distribution. We computed brain state transition cost with various *T* and observed that the results did not change qualitatively. Thus, we here show the results with *T* =1, i.e., the next time frame (see Supporting information S6 for results when *T* > 1). A probability distribution for each task was computed as an empirical probability distribution by counting the number of the occurrences of the coarse-grained states in each task time-series data. We estimated joint probability distribution of the resting state for two consecutive frames by counting transition pairs of the coarse-grained brain states with trajectory bootstrapping (see Methods for more details). From the joint probability distribution, we obtained the transition probability matrix of the resting state. Using these probability distributions and the transition probability matrix of the resting state, we calculated brain state transition cost represented by the minimized KL divergence (Eq. (4)). For instance, when we computed the transition cost from gambling task to motor task, we set the initial distribution, *π*, to be the empirical probability distribution obtained from gambling task data and the target distribution, *π*′, to be the empirical probability distribution obtained from motor task data. Here, the probability distribution of the uncontrolled path, *Q_ij_* = *q*(*X*_0_ = *i, X_T_* = *j*), is computed by the product of the initial probability distribution *π_i_* and the transition probability distribution of the resting state *Q_j|i_* = *q*(*X_T_* = *j |X*_0_ = *i*), *Q_ij_* = *Q_j|i_π_i_*. With *π*, *π*′, and *Q*, we can determine the optimally controlled path, *P*\*, and then compute the transition cost as 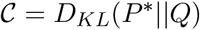.

We began by testing whether transition cost from rest to a more difficult task is larger. For this purpose, we quantified the transition cost from the distribution at rest to those during 0-back (easier) and 2-back (more difficult) tasks in the working memory (WM) task data. We chose the WM task because the WM task data are the only task data in HCP, wherein subjects perform tasks with objectively different levels of task difficulty. As shown in Fig. 3, we found that the transition cost to a 2-back task is larger than that to a 0-back task. This result suggests that our cost metric may capture the level of task difficulty from fMRI data.

**Figure 3:**
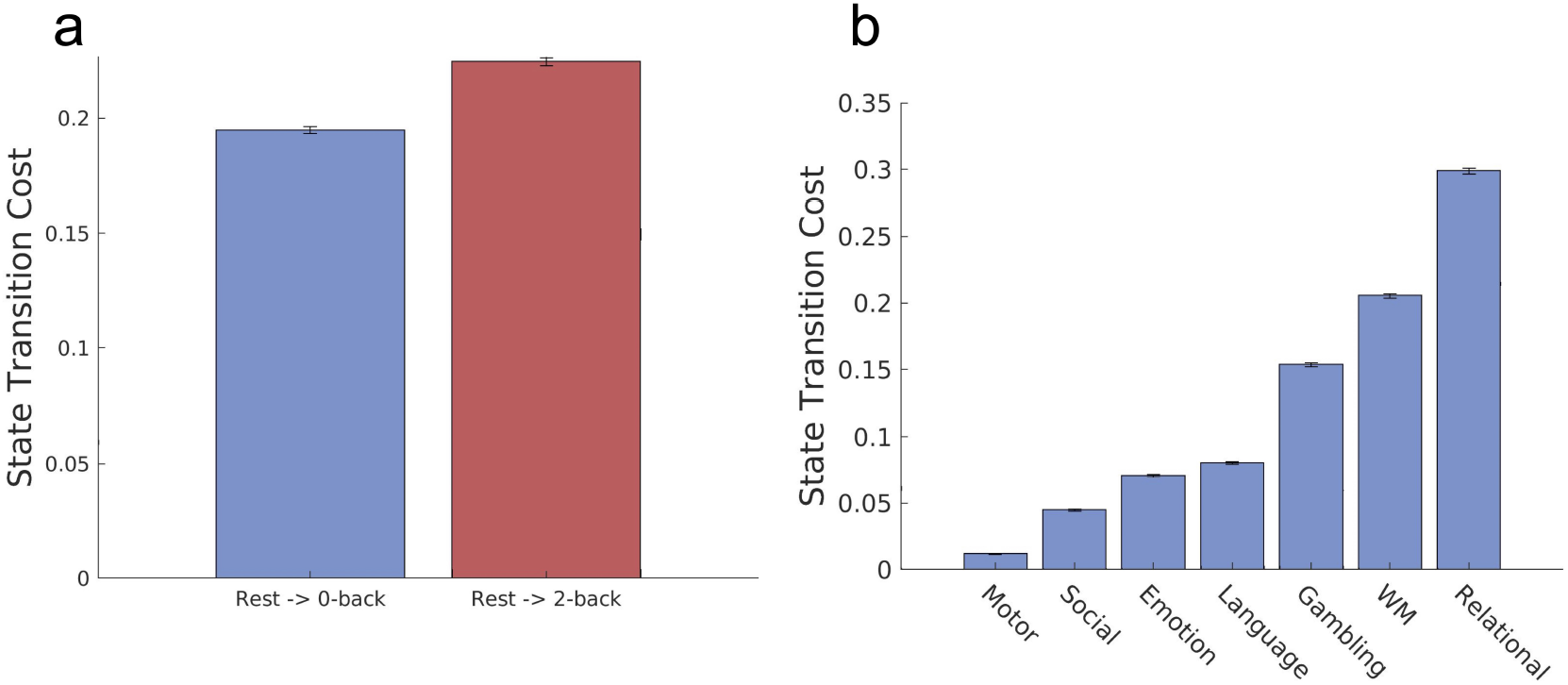
Brain state transition cost from the resting state to tasks. We computed the transition cost from the rest to cognitive tasks in the HCP data, as the minimum Kullback-Leibler divergence from the optimally controlled path to the baseline uncontrolled path. (A) The transition cost from the 2-back task (more difficult) is larger than the transition cost to 0-back task (easier) (one-sided t-test, *p* ≪ 0.001, *t* > 60, *df* = 198). (B) Transition cost from the rest to the seven cognitive tasks in the HCP dataset. Values are averaged over 100 bootstrapping trajectories and error bars indicate one standard deviation estimated with trajectory bootstrapping.

### Brain state transition cost to multiple tasks

To further check the behavior of the proposed metric for transition cost, we then computed brain state transition cost to multiple task distributions in the HCP dataset (emotion, gambling, language, motor, relational, social, and working memory). Note that, unlike the working memory tasks, we cannot objectively compare their task difficulties since these tasks are qualitatively different. Thus, the analysis here is exploratory without any prior hypothesis.

We found that the degree of transition cost to the seven cognitive tasks is significantly different. Fig. 3b shows the rank order of transition costs in the seven cognitive tasks. Notably, transition cost to a motor task was smallest among the seven tasks, whereas the transition cost to a relational task was the largest (see Discussion).

### Asymmetry of brain state transition cost

We then investigated whether state transition cost between tasks with different task difficulty was asymmetric. We hypothesized that it would require a larger transition cost to switch from an easier task to a more difficult task. To test this hypothesis, we computed the transition cost between 0-back and 2-back tasks in the working memory task. As shown in Fig. 4a, we found that the transition cost from a 0-back task to a 2-back task was larger than the cost accompanying the reverse direction, which agreed with our hypothesis (one-sided t-test, *p* ≪ 0.001, *t* > 80, *df* = 198). Note that the asymmetry of brain state transition cost does not result from the asymmetry of KL-divergence because the cost is not solely computed from the endpoint distributions but with underlying transition probability.

**Figure 4:**
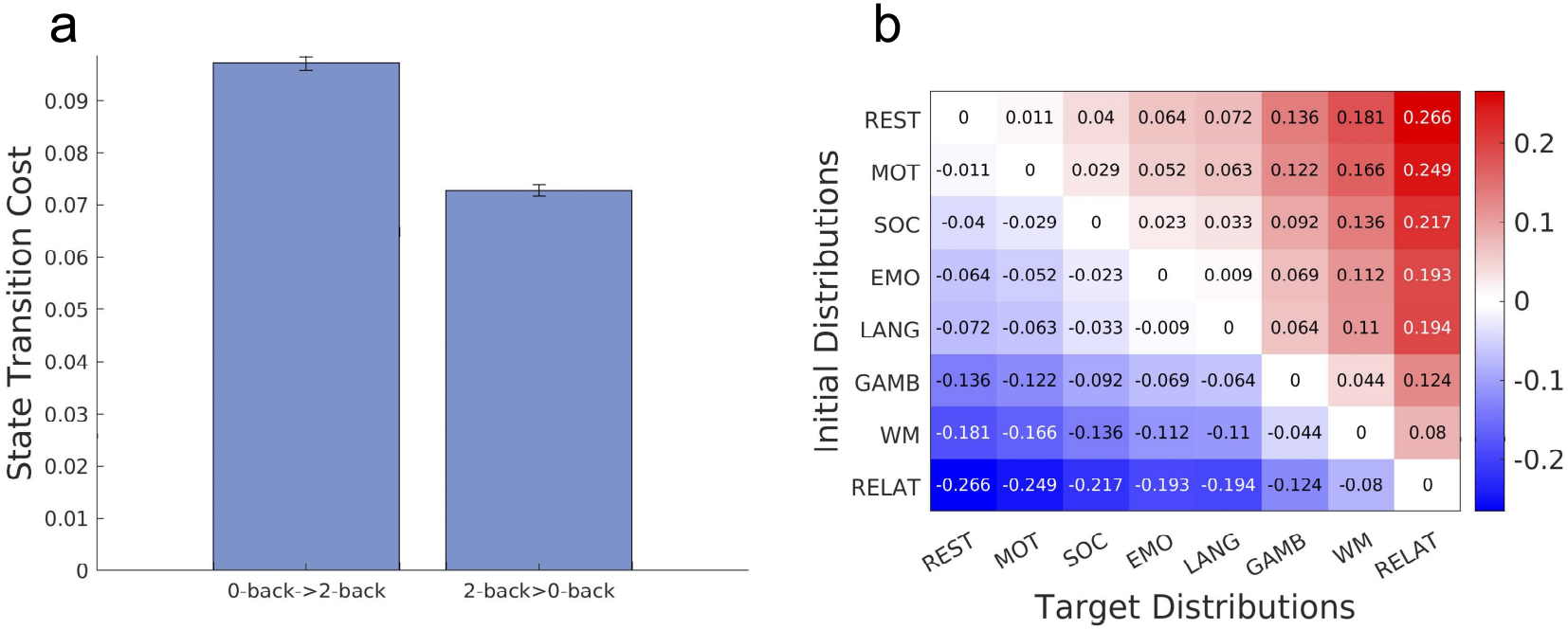
Brain state transition cost between task states. (a) Brain state transition cost between the 0-back and 2-back tasks in the working memory task. Values are averaged over 100 bootstrapping trajectories and error bars indicate one standard deviation estimated using trajectory bootstrapping. (b) Asymmetry of brain state transition costs for the rest and seven cognitive tasks. Each element in the matrix represents a difference in transition cost between tasks.

Finally, we examined whether the asymmetric property of brain state transition cost would be observed in other tasks whose task difficulties cannot be objectively compared. Here, we checked if the following relationship would hold for all the pairs of tasks (note that here we regarded the rest as a task):

If the transition cost from rest to task A is larger than that from rest to task B, then the transition cost from task B to task A is larger than that in the reverse direction,

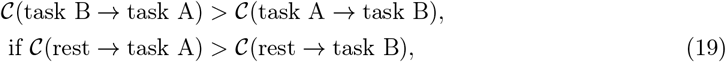

where 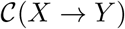 represents the brain state transition cost from X to Y, which is quantified by the KL divergence. To evaluate the relationship, we calculated the difference in transition cost between every pair of tasks, which is obtained as follows.

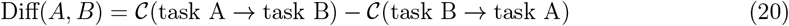

The result is summarized in the matrix in Fig. 4b, wherein entries (tasks) are arranged in ascending order by transition cost from rest, i.e., the first row (column) corresponds to the task with the smallest transition cost from rest and the last row (column) corresponds to the task with the largest transition cost from rest. The (*i, j*) entry of the matrix represents 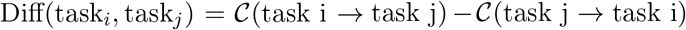. We observed that every entry in the upper (lower) triangular parts was positive (negative). This means that the relationship represented in Eq. 19 holds for every pair of tasks in the dataset. That is, the transition cost is asymmetric between tasks with different degrees of transition cost.

## Discussion

In this study, we propose a novel framework for quantifying brain state transition cost in stochastic neural systems by framing brain state transition as Schrödinger’s bridge problem (SBP). This framework resolves the problem of previous methods that cannot take account of the inherent stochastisity of neural systems^20,41^ while still utilizing principled control-theoretic approaches. Under this framework, we assumed that the brain follows the resting state activity as the baseline uncontrolled dynamics, and transitions to other distributions of brain state by modulating the baseline dynamics. Transition cost is measured as the minimum KL divergence from the uncontrolled path to the controlled path with the fixed endpoint probability distributions. We tested the utility of our framework by applying it to fMRI data from the Human Connectome Project release. The results indicated that the transition cost metric proposed in our framework may be useful for elucidating the characteristics of transition cost between tasks with different task difficulties.

### Correspondence between transition cost and cognitive demands

In the present study, we aimed to examine the relationship between the degree of brain state transition cost and task difficulty as proof of concept. We refer to task difficulty as objectively quantifiable task difficulty (e.g. 0-back and 2-back) only, not as a subjectively experienced task difficulty, which could vary among subjects. As for the objective task difficulty, we observed that the transition cost to a 2-back task (a more difficult task) is larger than that to a 0-back task (an easier task). Further studies using different types of tasks with various levels of difficulty are needed to determine the generality of this result.

On the other hand, we did not deal with subjectively experienced task difficulty or cognitive demands, as the dataset does not contain subjective reporting on the cognitive demand of each task. Nevertheless, we quantified the transition cost to the seven qualitatively different tasks, whose task difficulty or cognitive demand cannot be objectively quantified. Although it is unclear whether the observed order of transition cost correlates with subjective cognitive demand (Fig. 3b), one may at least consider it reasonable that transition cost to a motor task is substantially smaller than that to a relational task. This is because performing a motor task only requires subjects to move a part of their body (e.g., right hand or tongue) whereas performing a relational task requires processing multiple visual stimuli and reasoning their relationships, which appears significantly more demanding than performing a motor task. Although we could not further examine if the degree of transition cost correlates with the degree of cognitive demand, investigating this relation in more detail would be an interesting future work. We expect that while there may be a rough correlation between transition cost and cognitive demand, there can never be one-to-one correspondence, as many factors affect subjective evaluation of cognitive demand^42–45^. It would be intriguing to investigate the difference in transition cost and cognitive demand and in what cases these behave similarly or differently. The direction of study proposed in the present study might be an important step toward bridging cognitive demand and brain activity.

### Relation to previous theoretical work

In the present study, we considered the brain dynamics as a discrete stochastic process by coarse-graining brain activity patterns, which reduced the computational cost. On the other hand, previous studies using network control theoretic framework^15^ employed a linear continuous process. Our framework can also be extended to a linear continuous stochastic process because Schrödinger bridge problem is not limited to a discrete process but has also been studied in continuous settings as well^26–28^. We therefore expected that we could directly fit the high dimensional neural recording data with a continuous model (e.g. stochastic differential equations) as recently implemented^46^ and carry out a similar analysis as the present study. Developing a continuous version of the present framework may allow us to gain more insights into brain state transition cost.

Both the discrete stochastic process (model-free dynamics) utilized in the present study and linear dynamical models^15,29^ possess pros and cons for the application in the analysis of neural activity. In linear dynamical models, because control input is explicitly modeled, the biophysical meaning of control input is clear. By taking advantage of the high interpretability of control input, linear dynamical models, for example, can provide insights into the contribution of each brain region to the control of the whole system^15^. However, linear dynamical models are not suitable for the analysis of neural activity that is highly nonlinear. The discrete stochastic process used in the present study can be applied to nonlinear neural activity although the biophysical interpretation of control input is unclear. It is imperative to select an approach that fits the purpose of the study and the property of the data.By choosing appropriate models, we can compute brain state transition cost in various types of data (e.g., fMRI, EEG, ECoG, etc).

Similarly to our framework, some recent works have utilized information theoretic measures to quantify cognitive costs^47^ and connectivity changes between brain states^48^. While these studies compute only distance or divergence between the two distributions of brain state, our framework takes account of the underlying baseline activity of the brain. We employ this approach because including this baseline spontaneous activity provides a more accurate transition cost measure from the viewpoint of dynamical system theory.

### Physical interpretation of the brain state transition cost

The KL divergence-based control cost proposed in this study may seem to be a distant concept from the conventional control cost, the time integral of squared input^16^, in a linear deterministic model. However, it was shown in the previous study that the KL divergence cost in a stochastic linear model is analytically computed as the expectation of the time integral of squared input^25,29^. In this sense, the KL divergence cost is tightly connected to the conventional control cost in a linear system.

The brain state transition cost proposed in the present study has a clear information theoretic meaning, i.e., the KL divergence between the optimally controlled path and the uncontrolled path. However, an explicit physical interpretation has yet to be elucidated. A natural choice of control cost from the viewpoint of physics would be the work needed for realizing a controlled path from an initial distribution to a target distribution^49,50^. Minimizing the work is equivalent to minimizing entropy production (work dissipation)^49,50^. Previous works investigated the optimal control minimizing the entropy production (or equivalently the work) and showed that the entropy production is lower bounded by the square of the Wasserstein distance^50,51^. Interestingly, the entropy production is given by the Kullback-Leibler divergence between two probabilities of forward and backward processes^52^. Thus, one may consider that there should be some connections between Schrödinger bridge type information theoretic control cost proposed in this study and the physical cost, work or entropy production. It would be an interesting future work to clarify the relationship between these different types of control costs.

### Brain state transition and reconfiguration of functional connectivity

The brain state transition cost computed in our framework may be related to the reconfiguration of functional connectivity between tasks. Numerous functional neuroimaging studies have reported the alterations in functional connectivity from the resting state connectivity during task performance^53–57^. In our present study, we showed that transitioning to more difficult tasks carries a larger transition cost. This seems to be consistent with Kitzbichler’s work, which demonstrated that larger cognitive demand induces a more global alteration in brain activity^31^. It may be the case that our framework captures the cost associated with the degree of reorganization of functional connectivity between different tasks.

### Implications to task switching

The Schrödinger’s bridge-based framework we proposed in this study may provide a new perspective for studying task switching from brain activity data. One of the most important and replicated findings in the task switching paradigm is the observation of switch cost^30^: switching to a new task takes a higher reaction time and error rate than repeating the same task. Various hypotheses have been proposed to explain the source of switch cost^58,59^, including reconfiguration of the mental set for performing tasks. However, few studies have quantified the switch cost from brain activity. A recent work suggests that task switching involves the reconfiguration of brain-wide functional connectivity^60^. Our framework may be used as a quantitative method for measuring switch cost. Subsequent investigations should study the relationship between switch cost and brain state transition cost by measuring brain activity while a subject is performing a task-switching experiment.

### Comparison between the optimally controlled path and the empirical path

We investigated only the computed optimally controlled path, not an empirical path due to the limitation of fMRI recording. To quantify the efficiency of brain state transition, it would be interesting to compare empirical and optimal paths. This may provide insight into individual differences in the performance of task switching. However, the fMRI data from Human Connectome Project does not include recordings in which subjects perform and switch between multiple tasks, and we were therefore unable to compute an empirical transition path between initial and target distributions of brain states. Even if the dataset contained such data, fMRI would not capture rapid transitions between tasks because the time resolution of the fMRI data is not sufficiently high (TR = 0.72 second in the HCP data set). Computing empirical transition paths will require the use of recording data with better temporal resolution, such as EEG, MEG, ECoG, etc. In this regard, our theoretical framework is applicable to other types of recording data beside fMRI.

## Methods

### fMRI data acquisition and preprocessing

The 3T functional magnetic resonance imaging (fMRI) data of 937 subjects were obtained from the Washington University-Minnesota Consortium Human Connectome Project (HCP)^32^. Every subject provided a written informed consent to the Human Connectome Project consortium, following the HCP protocol. We used minimally preprocessed fMRI data at resting state and seven cognitive task states (emotion, gambling, language, motor, relational, social, and working memory). We selected these 937 subjects as they contain complete data for all the tasks. We then performed denoising by estimating nuisance regressors and subtracting them from the signal at every vertex^61^. For this, we used 36 nuisance regressors and spike regressors introduced in a previous study, consisting of (1-6) six motion parameters, (7) a white matter time series, (8) a cerebrospinal fluid time series, (9) a global signal time series, (10-18) temporal derivatives of (1-9), and (19-36) quadratic terms for (1-18). Following a previous study^61^, the spike regressors were computed with 1.5 mm movement as a spike identification threshold. After regressing these nuisance time courses, we also applied a band-pass filter (0.01-0.69 Hz) to the data, in which the upper bound of the filter corresponds to the Nyquist frequency of the time-series. We then applied a parcellation proposed in^40^ to divide the cortex into 100 brain regions, which reduced the complexity of the following analysis.

### Clustering BOLD signals

In order to model brain dynamics as a discrete stochastic process, we coarse-grained brain activity patterns using the *k*-means clustering algorithm. While there are numerous unsupervised clustering algorithms, we chose the *k*-means clustering due to its effective fit with the dynamics of neural activities^17^. We used cosine similarity as a distance measure between centroids of clusters, which is commonly used in high dimensional data. As described in the previous studies^17,33^, we concatenated preprocessed BOLD signals of all the subjects during the resting state and seven cognitive tasks. We obtained a *M* × *N* matrix, *X*, where *M* is the number of parcels (100), and *N* is the number of task types times the number of time frames times the number of subjects. In order to prevent the variability in data size across tasks from affecting the clustering results, we used the same number of time frames for each task data. We used a different number of time frames depending on whether we divided the working memory task data into 0-back and 2-back tasks or not. When we divided the working memory task data (Figs. 3a and 4a), we obtain 148 time frames that included either 0-back or 2-back task blocks. Accordingly, we only used the first 148 time frames in the other tasks. When we did not divide the working memory task data (Figs. 3b and 4b), we used only the first 176 discrete measurements in each task since the emotion task - the task with the shortest measurement - was recorded for 176 time frames.

We determined the number of clusters using a similar procedure to that in previous studies^17,33^. We first computed the percent variance explained by the number of clusters varying from *k* = 2 to *k* = 12. We observed that the explained variance plateaued around 75% after *k* = 5 (S4a in Supporting information). We then examined whether all the coarse-grained states would appear in every subject during each task session (S4b). We found that when we set the number of clusters to be greater than *k* = 8, some coarse-grained states did not appear in the data of some subjects. For these two reasons, we selected the number of clusters to be 8. While we chose *k* = 8, we were able to reproduce the major results with *k* > 4 (S1-S3), which indicates the robustness of our results regardless of the number of clusters.

### Estimating transition probabilities and probability distributions of coarse-grained states using trajectory bootstrapping

We are limited by the finite length of the time-series data for estimating brain state transition cost, which are calculated using probability distributions and transition probabilities. To ensure the accuracy of estimated quantities, we applied trajectory bootstrapping^33,62^ to calculate error bars on the estimated quantities. After we classified brain activity data with *k*-means clustering, we estimated the joint probability distribution matrix, *M_ij_* = *q*(*X_τ_* = *i,X*_*τ*+1_ = *j*), of coarse-grained states at time *t* = *τ* and *t* = *τ* + 1 for the resting state. To obtain the matrix, we first created a list of transitions in concatenated time-series data of the resting state, in accordance with previous work^33^

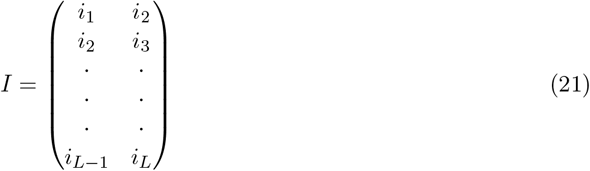

where *i_l_* is the coarse-grained state at *l*-th frame of the time-series, and *L* is the length of the concatenated time-series (*L* = the number of time frames × the number of subjects). We sampled a pair of transitions from the list for *L* times to fill in the matrix, *M*, and normalized it to be a joint probability matrix. Although the transition list is a concatenated time-series across subjects, we excluded pairs of transitions that took place across subjects; we only sampled pairs within the same subject. By normalizing each row of the matrix, *M*, to be 1, we constructed a transition probability matrix for the resting state. Similarly, we computed probability distributions of coarse-grained brain state using trajectory bootstrapping. From the concatenated time-series data of each task, we sampled a coarse-grained brain state *L* times. We counted the number of the occurrences of each coarse-grained brain states and normalized it to 1 to obtain the probability distribution for each task. While we calculated a probability distribution for each task including the rest, we computed transition probability only for the resting state for obtaining the uncontrolled path, *Q*. We followed this process 100 times and computed the error bars on the estimated quantities in this study using the 100 bootstrap trajectories.

## Acknowledgments

This work was supported by JST Moonshot R&D Grant Number JPMJMS2012, JST CREST Grant Number JPMJCR1864, and JSPS KAKENHI Grant Number 18H02713, Japan.

## Code Availability

The code for computing the transition cost based on optimal control for stochastic systems is available at https://github.com/oizumi-lab/SB_toolbox.

## Supporting Information

Supporting information includes the following supplementary figures. S1: The robustness with different numbers of clusters. S2: The order of the degrees of transition costs from the resting state. S3: The asymmetry of brain transition cost. S4: Criteria for determining the number of clusters for k-means clustering algorithm. S5: The figures of brain maps. S6: Brain state transition cost when the time horizon, *T*, is set *T* > 1.

---

**Figure S1:**
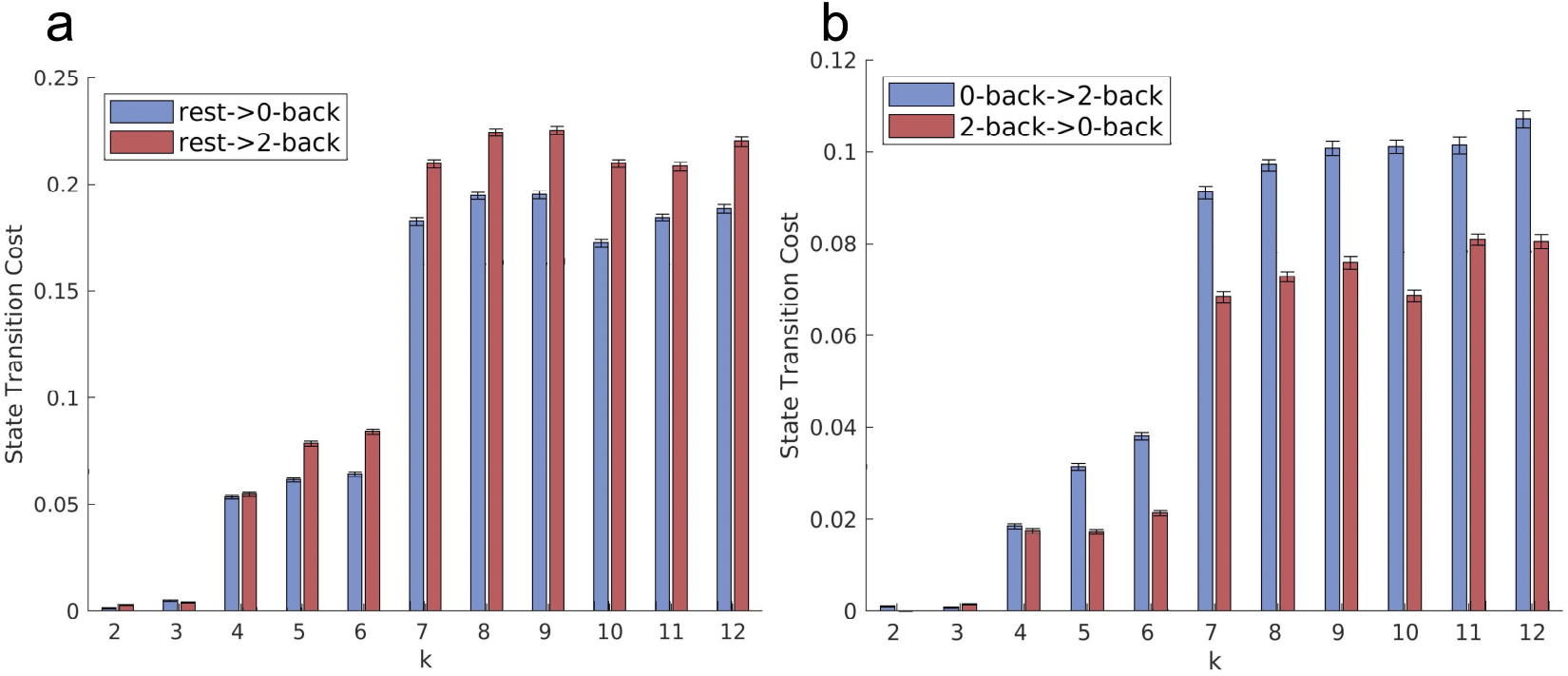
The robustness with different numbers of clusters. **(a)** Brain state transition cost from the resting state to 0-back and 2-back tasks. For *k* ≥ 4, the cost for transitioning to 2-back task is larger than that of 0-back task, which is consistent with the result in the manuscript with *k* = 8 (Figure 3a). **(b)** Brain state transition cost between 0-back task and 2-back task. For *k* ≥ 4, the cost for transitioning from 0-back task to2-back task is larger than that of the opposite direction. This is in line with the result in the main text with *k* = 8 (Figure 4a). The results in (a) and (b) indicate the robustness of the methodology used in the manuscript with respect to the number of clusters for *k*-means clustering.

**Figure S2:**
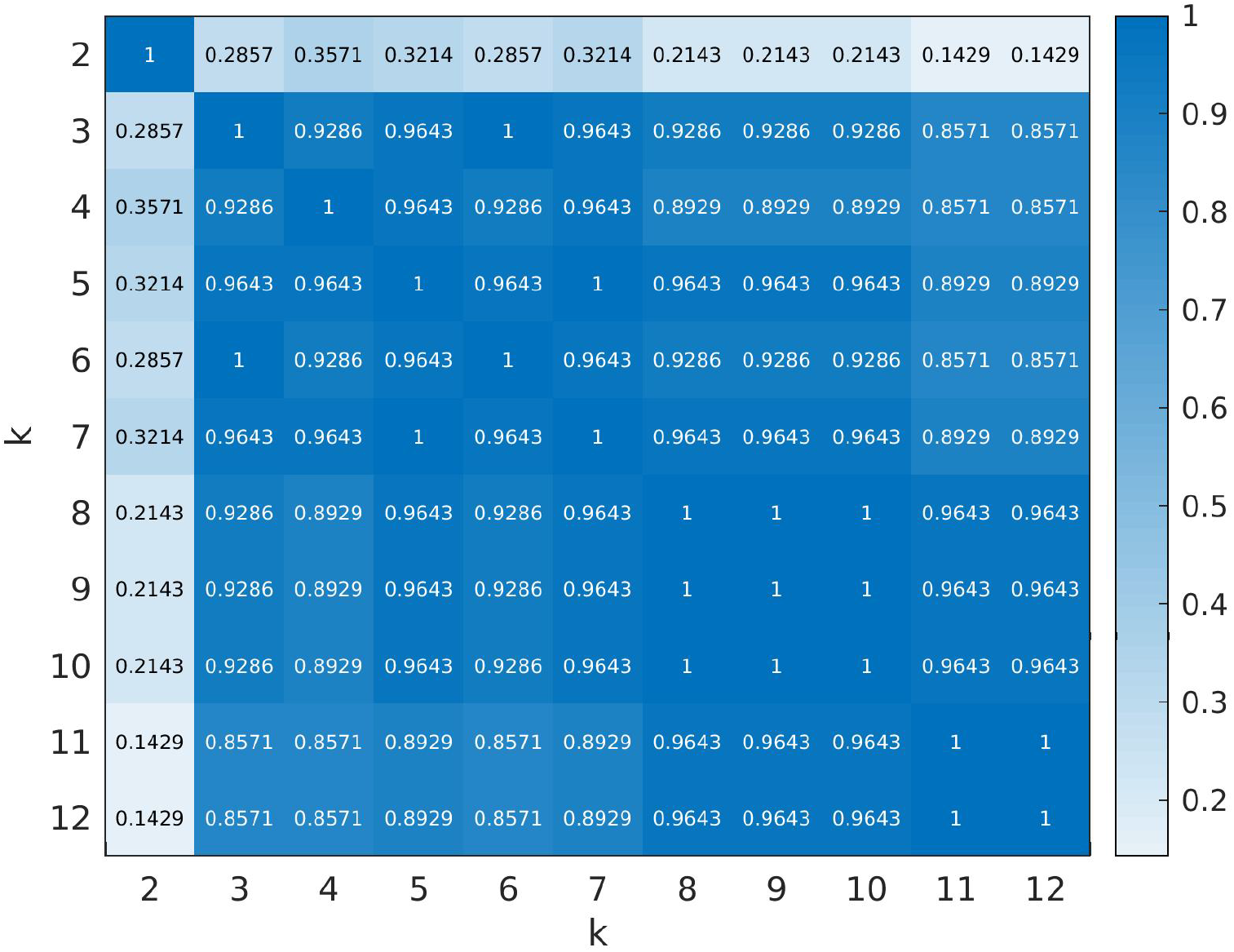
The order of the magnitudes of transition cost from the resting state. Spearman’s rank correlation coefficients between the order of the magnitudes of transition cost from the resting state for different numbers of clusters. For 2 ≤ *k* ≤ 12, we computed the Spearman’s rank correlation coefficient between the order of the magnitude of transition cost from the resting state to the seven cognitive tasks (emotion, gambling, language, motor, relational, social, and working memory) for different number of clusters. The figure shows that the order is highly consistent for *k* ≥ 3, which demonstrates the robustness of the result in the manuscript (Figure 3b).

**Figure S3:**
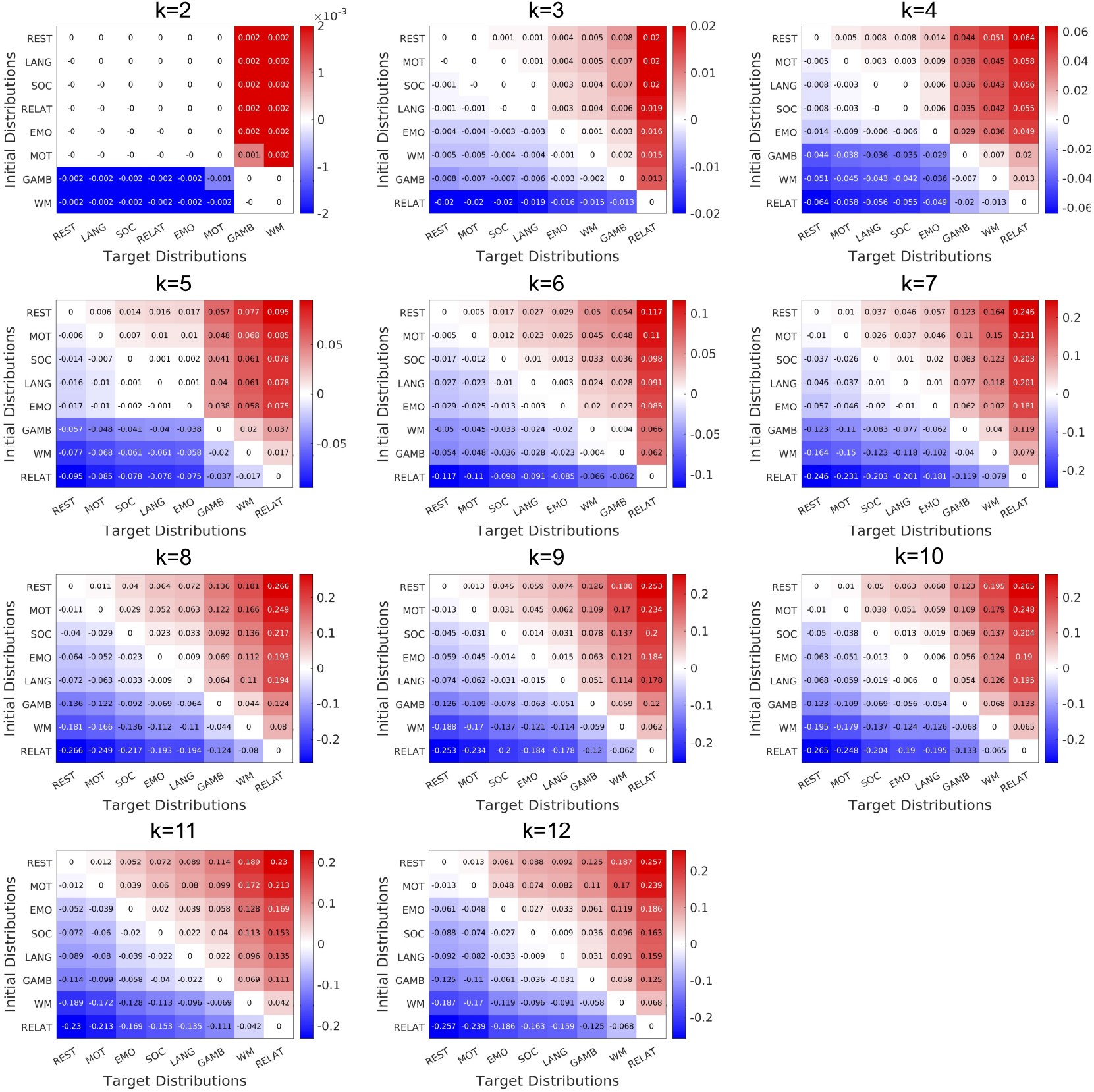
The asymmetry of brain state transition cost. Asymmetry of brain state transition cost for different number of clusters. We tested whether the asymmetric relationship for transition costs between eight tasks including the resting state (emotion, gambling, language, motor, relational, social, and working memory) (see Results for the definition) may hold for *k* ≠ 8. As shown in the figure, we observed that the asymmetry of brain state transition cost between all the pairs of tasks for 2 ≤ *k* ≤ 12.

**Figure S4:**
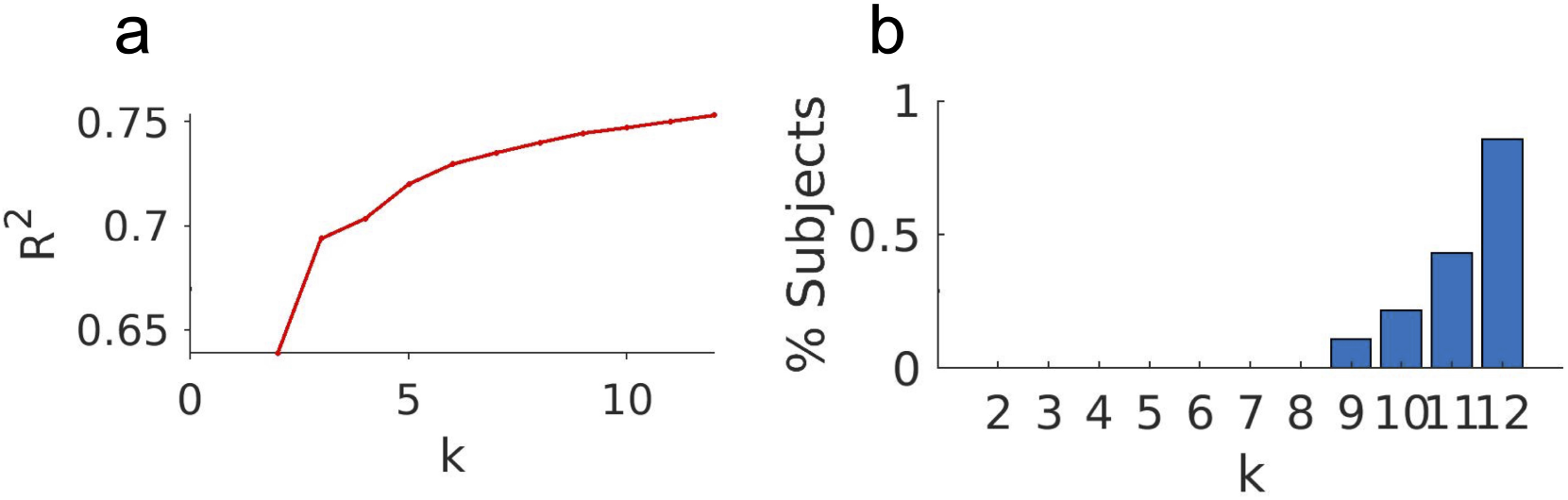
Criteria for determining the number of clusters for k-means clustering algorithm. (a) The percentage of variance explained by the number of clusters varying from *k* = 2 to *k* = 12. We observed that the explained variance plateaued around 75% after *k* = 5. (b) The percentage of subjects missing a cluster (brain state) from one task session. For *k* > 8, we observed that some clusters were missing from some subjects. Based on the results in (a) and (b), we set the number of clusters to be *k* = 8 for the analysis in the manuscript. However, as shown in the S1-S3, our main results are robust for 4 ≤ *k* ≤ 12.

**Figure S5:**
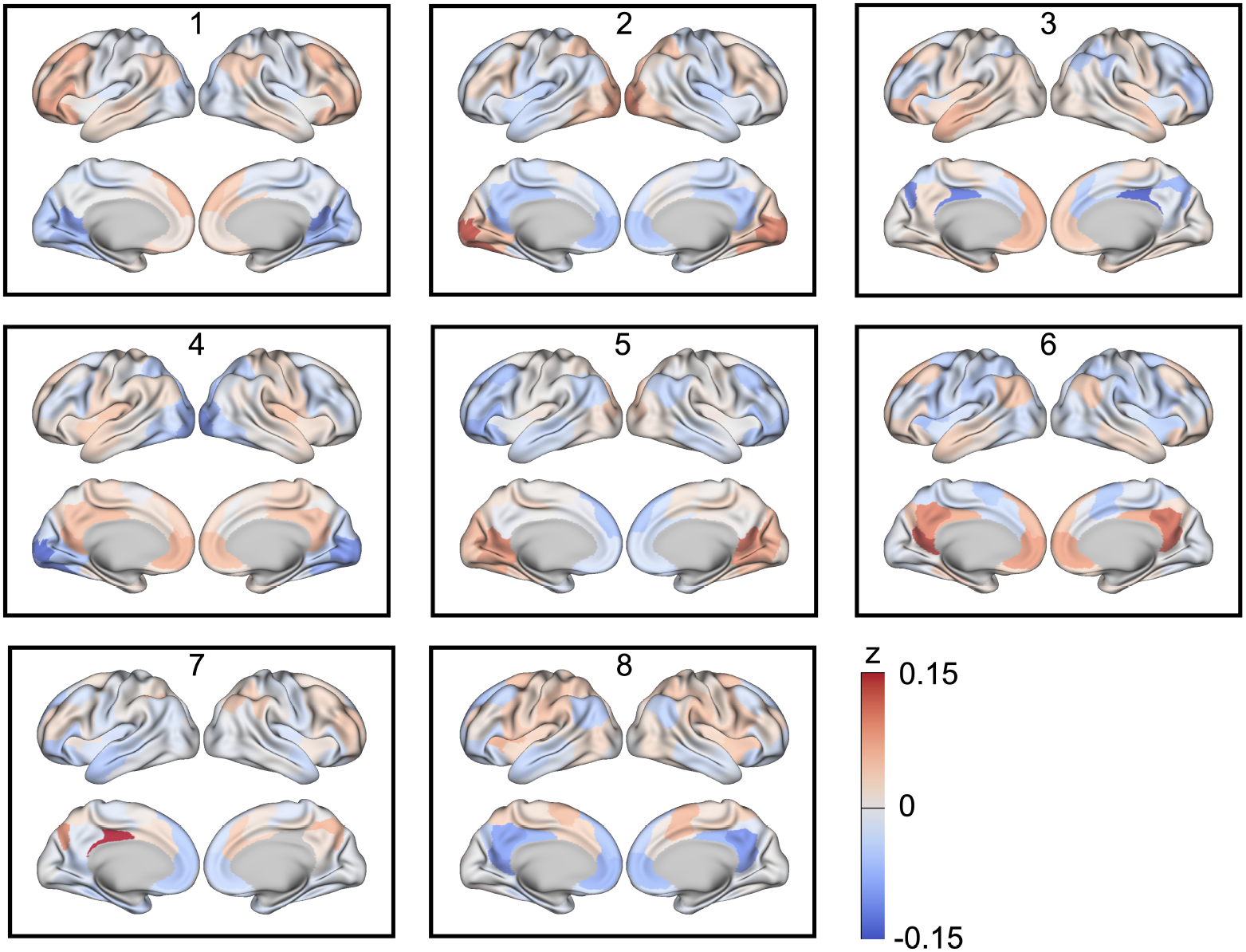
Figures of brain maps. The z-scored activities of eight brain states defined by the centroids of eight clusters determined by *k*-means clustering algorithm. Brain regions with high-amplitude activities are colored red and those with low-amplitude activities are colored blue.

**Figure S6:**
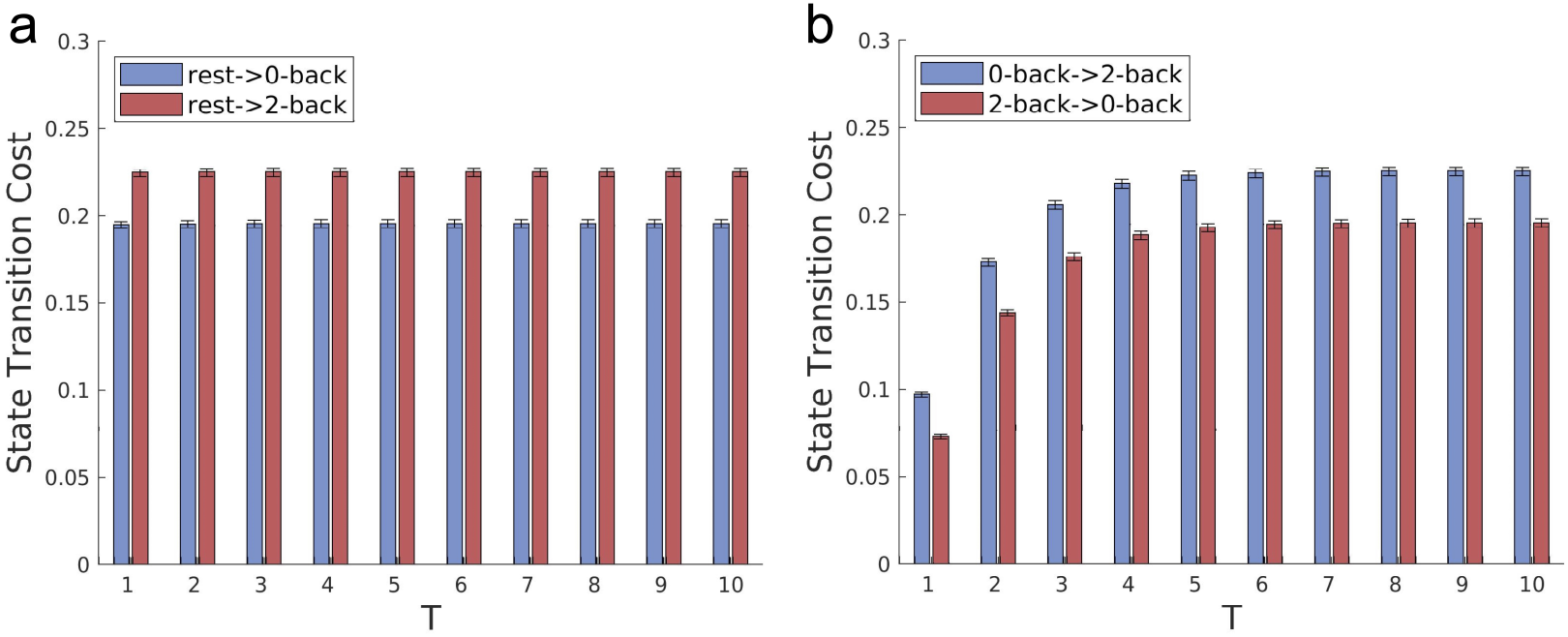
The reproducibility of the results with *T* > 1. Brain state transition cost when the time horizon, *T*, is set *T* > 1. **(a)** Brain state transition cost from the resting state to 0-back and 2-back tasks with *T* ranging from 1 to 10. Increasing *T* does not affect the degree of the transition cost because the initial distribution is set to be the probability distribution of the resting state, which is the stationary distribution of the transition probability of the uncontrolled path. **(b)** Brain state transition cost between 0-back and 2-back tasks. Increasing *T* makes the cost larger up to *T* = 4, but the degree of cost stays the same for *T* > 5, as the endpoint distribution of the uncontrolled path, *q*(*X_T_*) converges to the stationary distribution of the resting state transition probability, as *T* becomes larger.

